# Hypothesis-driven probabilistic modelling enables a principled perspective of genomic compartments

**DOI:** 10.1101/2022.10.01.510432

**Authors:** Hagai Kariti, Tal Feld, Noam Kaplan

## Abstract

The Hi-C method has revolutionized the study of genome organization, yet interpretation of Hi-C interaction frequency maps remains a major challenge. Genomic compartments are a checkered Hi-C interaction pattern suggested to represent the partitioning of the genome into two self-interacting states associated with active and inactive chromatin. Based on a few elementary mechanistic assumptions, we derive a generative probabilistic model of genomic compartments, called deGeco. Testing our model, we find it can explain observed Hi-C interaction maps in a highly robust manner, allowing accurate inference of interaction probability maps from extremely sparse data without any training of parameters. Taking advantage of the interpretability of the model parameters, we then test hypotheses regarding the nature of genomic compartments. We find clear evidence of multiple states, and that these states self-interact with different affinities. We also find that the interaction rules of chromatin states differ considerably within and between chromosomes. Inspecting the molecular underpinnings of a four-state model, we show that a simple classifier can use histone marks to predict the underlying states with 87% accuracy. Finally, we observe instances of mixed-state loci and analyze these loci in single-cell Hi-C maps, finding that mixing of states occurs mainly at the population level.

## Introduction

In the past decade, genomic methods (1–7) have highlighted that the spatial organization of the genome is closely intertwined with a wide variety of physiological processes, including transcription (8–11), replication (12–14), sex chromosome inactivation (15–18), development (19–22), mitosis (23–27) and spermatogenesis (28–31). One of the most popular of these methods is Hi-C (1, 32, 33), a genome-wide assay which uses proximity ligation to measure interaction frequencies between every pair of loci in the genome within a cross-linked population of cells. Since genome structure is highly stochastic, the resulting Hi-C interaction frequency matrix represents not a single structure but a distribution of structures. Structural features of the genome which are consistent in the cell population, constrain this distribution and manifest as patterns in the interaction map. The identification of such patterns, their interpretation, and their molecular specification remain outstanding challenges in the field of genome organization.

Hi-C interaction patterns appear across many scales. At the whole-chromosome level, intrachromosomal (cis) interactions are much more frequent than interchromosomal (trans) interactions, due to a combination of the physical separation of chromosomes into territories and the stochastic positioning of territories within the nucleus (34, 35). Within chromosomes, interaction frequency between pairs of loci tends to decrease – on average – as a function of their genomic distance, often following a power-law decay (36–38). Although these large-scale structures are not locus-specific, they can provide useful information on general polymer properties and have also separately been found useful in genome assembly-related applications (39–44). At the multi-Mb scale, genomic compartments are a checkered interaction pattern found both in and between chromosomes (1, 45, 46). Genomic compartments were initially suggested to represent a partitioning of loci into two states, where loci of similar states interact more frequently than loci of different states. At the sub-Mb scale, Topologically Associating Domains are patterns of genomic domains in which loci within a domain interact with each other more frequently than with loci outside the domain (46–48). Although both genomic compartments and TADs form secluded self-interaction domain-like structures in Hi-C maps, TADs are likely associated with a completely different mechanism, namely the action of cohesin-mediated loop extrusion which can be stopped at boundaries which often involve CTCF (49–53).

In this work, we focus on genomic compartments while also accounting for larger structures. Relative to TADs, for which a plethora of approaches have been developed to detect and model, the method used for identifying genomic compartments in the original Hi-C paper (1) is still the standard for detecting genomic compartments. This typically involves normalizing the interaction matrix by average interaction frequency per genomic distance and then applying Principal Component Analysis. The first eigenvector is then taken to represent the genomic compartment signal, where positive values are assigned to one compartment and negative values to the second compartment. Although widely used, this heuristic is intrinsically limited as it is not a model. Therefore, it is not an explanation of the observations, it does not explicitly test hypotheses about the underlying mechanisms, its parameters are not directly interpretable, and it is not generative or predictive. Additionally, this method suffers from technical limitations including insensitivity to scaling, an arbitrary threshold for partitioning of states, occasionally poor performance on noisy and sparse data, and an underlying implicit two-state assumption. In spite of these limitations, the PCA-based analysis has proved quite useful in providing a genomic track that heuristically quantifies genomic compartment signal. Most notably, comparing this track with other genomic features shows some correspondence between the interaction states (compartments) with known chromatin epigenetic states (1): Compartment A is usually euchromatic and tends to have higher GC content, higher gene density, and histone marks associated with active chromatin; Compartment B is usually heterochromatic and tends to have lower GC content and gene density, and is enriched in lamina-associated domains and repressive histone marks.

A range of computational techniques has been used to analyze and model genomic compartments. One set of approaches is the data-driven heuristic partitioning of loci to compartments (compartment “calling”), usually based on unsupervised learning, comparable to the common PCA method(1, 54): Rao et al. (45) used Gaussian HMM clustering interchromosomal interactions to define five subcompartments from human Hi-C maps, and observed characteristic histone marks; Yaffe et al. (55) used k-means clustering on interchromosomal contacts to identify three compartments; Nichols et al. (56) used k-means clustering to partition human Hi-C maps to four compartments, also matching different histone marks; Zheng et al. (57) used a probabilistic approach to calculate the CScore reflecting the likelihood of being associated with A or B states; and Rowley et al. (9) proposed an improved heuristic called the A-B index based on associating the interactions of high-resolution bins with a low-resolution genomic compartments track. Another set up approaches predict genomic compartment interaction patterns from one-dimensional features, usually with machine learning approaches: MEGABASE (58) uses a neural network to predict subcompartments from histone mark ChIP-seq data, and can be used with the MiChroM (59) physical model to directly predict the Hi-C matrix from histone modifications tracks; Rowley et al. (9) used a regression model to predict interactions from GRO-Seq and architectural protein binding site data in non-mammalian eukarya; Fortin et al. (60) use DNA methylation correlation matrices to predict A/B compartments; Nichols et al. (56) used a probabilistic model to predict human and drosophila Hi-C interactions from histone marks; SNIPER (61) uses a neural network to impute missing data in low-coverage Hi-C maps and classify loci to subcompartments; Esposito et al. (62) combine machine learning and polymer modeling to predict mammalian interaction maps from histone marks; and Orca (63) uses a neural network to predict regions of the Hi-C map from DNA sequence, with a tradeoff between resolution and size of the predicted region. Finally, a set of hypothesis-driven approaches based on polymer modelling have shown that multistate chromosomes can form interactions resembling genomic compartments: Jost et al. (64–67) used lattice-based block copolymer models and molecular dynamics to reproduce Hi-C maps form epigenetic states in various species; Mirny et al. (68, 69) used block copolymer models and Langevin dynamics to reproduce compartment patterns observed in Hi-C; and Nicodemi et al. (62, 70–72) used strings and binders polymer models to reproduce Hi-C maps at different scales based on a set of chromatin states.

Here we present deGeco, a generative probabilistic modelling approach to genomic compartments, which attempts to utilize some of the best properties of both hypothesis-driven and data-driven approaches. On one hand, we derive our model directly from a handful of explicit mechanistic assumptions. This makes the parameters of our model interpretable and makes the model suitable for testing biological hypotheses, in contrast to black-box machine learning approaches. On the other hand, the model is does not require polymer modelling and is data-driven, leveraging the data to infer the values of genome-wide biological parameters. After evaluating the performance and technical capabilities, we proceed to test several biological hypotheses regarding state-state interaction rules, interaction differences within and between chromosomes, the number of states, molecular underpinnings of states, and state mixing.

## Results

### Probabilistic model of genomic compartments

We propose a generative probabilistic model for genomic compartments based on the following four elementary assumptions on the mechanisms of the underlying system:

1. Every genomic locus in every cell is in one of *S* states. At the population level, every genomic locus is associated with a probability to be in each state.
2. Locus states are statistically independent.
3. Pairwise interaction probability decreases with genomic distance.
4. Pairwise interaction probability is specified by an affinity between the states of the respective loci.

These assumptions are sufficient to formulate the following model for pairwise interaction probability:

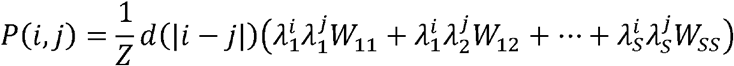

Where 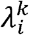 is the probability over the cell population that locus *i* is in the *k*-th state; *W*_*km*_ is a non-negative affinity between states *k* and *m*; *d*(|*i − j*|) is a distance-dependent function (we use |*i − j*|^*α*^ based on common polymer physics models); and *Z* is a normalization factor. Note that *W*_*km*_= *W*_*mk*_ since interaction is a symmetric property, and that from our first assumption we get 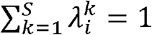. Also note that the model can be given a statistical mechanics interpretation, by using the Boltzmann distribution to convert state probabilities or interaction probabilities into energies of whole-genome configurations, under the assumption of thermal equilibrium.

We note that if we define a matrix Λ of locus state probabilities such that 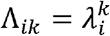 this model can be rewritten in matrix form (**Figure 1**):

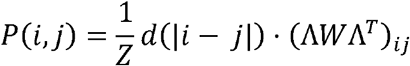

In this form, the model can be interpreted as performing a form of matrix factorization of the genomic compartments signal into the right-stochastic state locus probability matrix and the symmetric non-negative state-state affinity matrix. Accordingly, we call the model and its implementation deGeco (decomposition of genomic compartments). We note that if we set *S* = 2,*W* = *I*_2_ and *d*(|*i* − *j*|) to be a non-parametric function, the model resembles the approach of Zheng et al(57).

**Figure 1.**
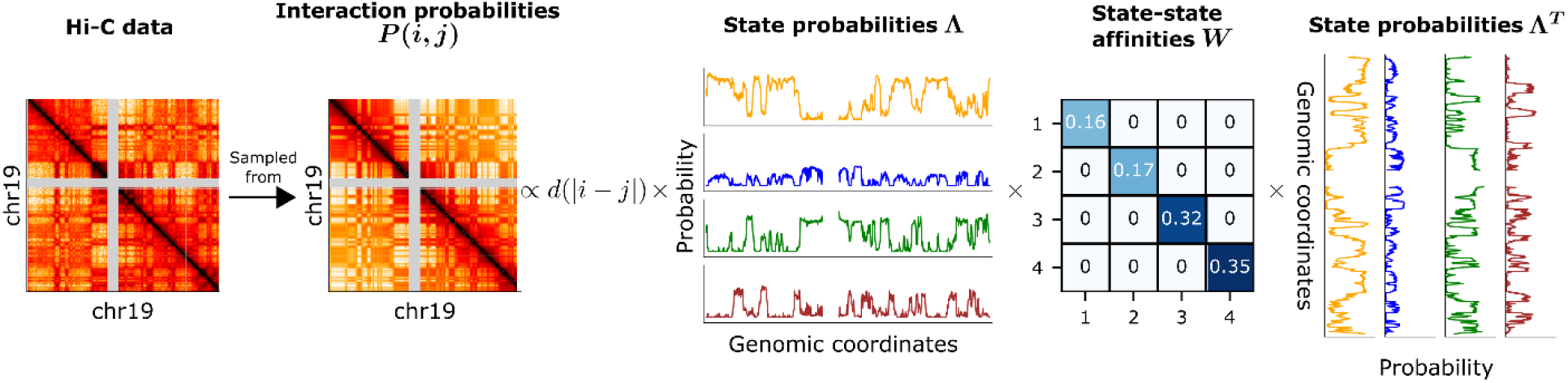
Overview of probabilistic model of genomic compartments. We assume that the Hi-C interaction frequency matrix is sampled from an underlying interaction probability matrix. Interaction probabilities result from two components: A distance-based interaction probability function and a state-based interaction probability component representing the genomic compartment signal. In the state-based interaction probability of two loci depends on the probability of each locus to be in each of the states (represented by matrix) and the affinities of these states to each other (represented by matrix). We show that the state-based interaction probability component is equivalent to a multiplication of these matrices.

As we cannot directly measure the interaction probabilities, we assume that the Hi-C experiment samples read-pairs independently from the underlying interaction probabilities. Thus, given a Hi-C interaction matrix, we can estimate the maximum likelihood parameters, and. Since the size of Hi-C matrices can be prohibitive, we developed a multiresolution fitting scheme using sparse data structures, which reduces the memory and CPU time requirements to get a good fit (see Methods). Alternatively, given parameter values, we can generate an interaction probability matrix or an Hi-C-like interaction frequency matrix sampled from these probabilities. Since the model parameters are interpretable and biologically meaningful, we can also simulate genomic perturbations and predict their effects on the Hi-C matrix.

### Explaining intrachromosomal genomic compartments with a two-state model

We first asked how well a two-state model can explain intrachromosomal (cis) interaction maps. We selected the deeply sequenced GM12878 interaction map of Rao et al.(45) at 50kb resolution, and fit the model separately to each chromosome (see Methods). Given the estimated parameters, we could calculate a predicted interaction probability matrix, and compare it to the Hi-C matrix. Since most of the variation in interaction maps can be explained by distance-dependent interaction, we calculated the Spearman correlation coefficient between the Hi-C and predicted matrices after first normalizing each of them by distance-dependent interaction. Importantly, the Hi-C matrix reflects a very sparse random sample from the true interaction probabilities, which sets an upper bound on the optimal possible correlation. Thus, we estimated for each chromosome the optimal possible correlation (see Methods) and compared this to the correlation achieved by the two-state model (**Figure 2A**). We find that the two-state model achieves a mean correlation of 0.54 (0.10 s.d.), and that this is on average 0.11 (0.04 s.d.) less than the optimal possible correlation. This correspondence is also apparent when visually comparing the data and the model (**Figure 2B, 2C**). Interestingly, we noticed that the model does not explain well TAD patterns (**Figure 2D**), supporting the notion that TADs are not simply small compartment domains but are rather due to a distinct mechanism (68, 73). Thus, a simple two-state model is sufficient to capture most of the explainable variation in the data.

**Figure 2.**
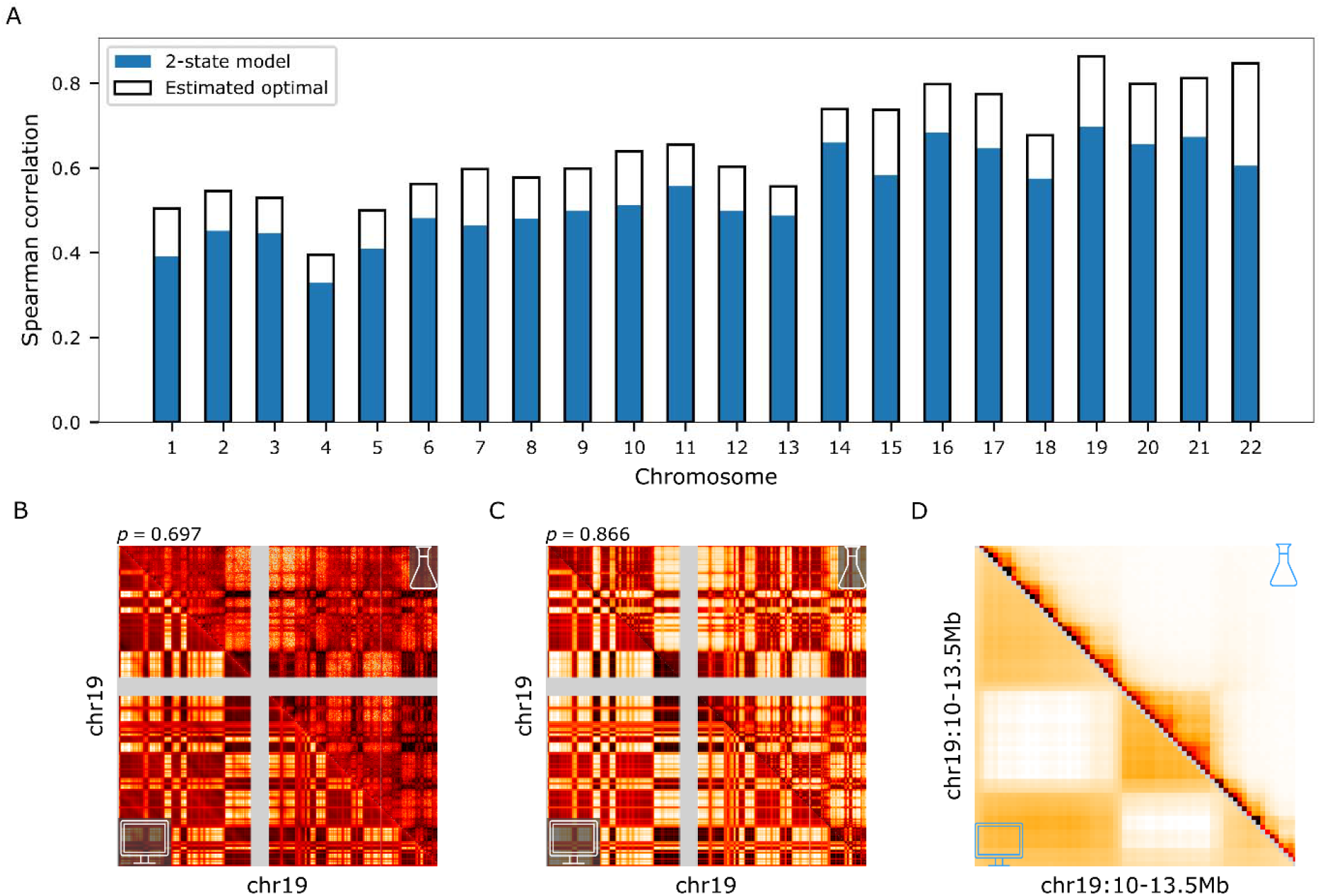
Two-state intrachromosomal (cis) model performance. The model was fitted to GM12878 Hi-C by Rao et al. (45) at 50kb resolution. (A) Distance-normalized Spearman correlation between the Hi-C interaction frequency matrix and the model’s inferred interaction probability matrix. The optimal possible correlation for the model at matching resolution and sequencing depth is shown as reference (see Methods for details). (B) Chromosome 19 Hi-C interaction frequencies (distance-normalized, upper triangle) versus the model-inferred interaction probabilities (distance-normalized, lower triangle). (C) Chromosome 19 Hi-C correlation matrix (distance-normalized, upper triangle) versus the model-inferred genomic compartments component (distance-normalized, lower triangle). (D) Closeup of a chromosome 19 3.5Mb region, showing Hi-C interaction frequencies (distance-normalized, upper triangle) versus the model-inferred interaction probabilities (distance-normalized, lower triangle). Genomic compartments appear in both the data and model, but TADs are apparent only in the data.

As the two-state model explains most the genomic compartments pattern, we next decided to take advantage of the interpretability of our model and turned to investigate the inferred parameters. Specifically, we wanted to verify that the inferred affinities are consistent across chromosomes, to check that the affinity between states is lower than the self-affinities of both states and to examine the hypothesis that the state self-affinities are equal. Examining the inferred affinity matrix across all chromosomes, we find the average affinity between states to be 0.0007 (0.002 s.d.), while the self-affinities of both states were approximately 0.5, with an average difference of 0.05.

### Recovering genomic compartments from sparse data

In Hi-C, read-pairs populate an interaction matrix whose size is the square of the number of bins. As a result, interaction maps are essentially always inadequately sampled and cost-restrictive sequencing depth directly affects the map resolution. To address this challenge, recent efforts have attempted to use black-box machine learning, trained on highly sampled Hi-C interaction maps, to enhance sparsely sampled interaction maps(74–77). We thus decided to evaluate the ability of our model to maintain its predictive performance using extremely shallow sequencing. To test the ability of our model to infer parameters and interaction probabilities on sparse data, we randomly down-sampled the GM12878 chromosome 19 interaction map at different sampling rates. We then applied the model to each random sample and compared the model’s predicted interaction probability with the original unsampled interaction frequency matrix across resolutions ranging from 500kb to 10kb-size bins, indicating how much performance decreases due to sequencing depth. We also compared the state probabilities inferred from each sample to the state probabilities that were inferred from the original data, to test the stability of the inferred parameters. Remarkably, we find only negligible deterioration in performance upon down-sampling (**Figure 3A, 3B**). Even when taking only 0.5% of the reads (equivalent to ∼20M reads genome-wide) at 10kb resolution, correlation decreases by only 0.02 and the average error in state probabilities is 0.07 (0.05 s.d.). Thus, our model can reconstruct high-quality interaction maps from extremely sparse interaction maps, simply based on mechanistic assumptions without any need for training.

**Figure 3.**
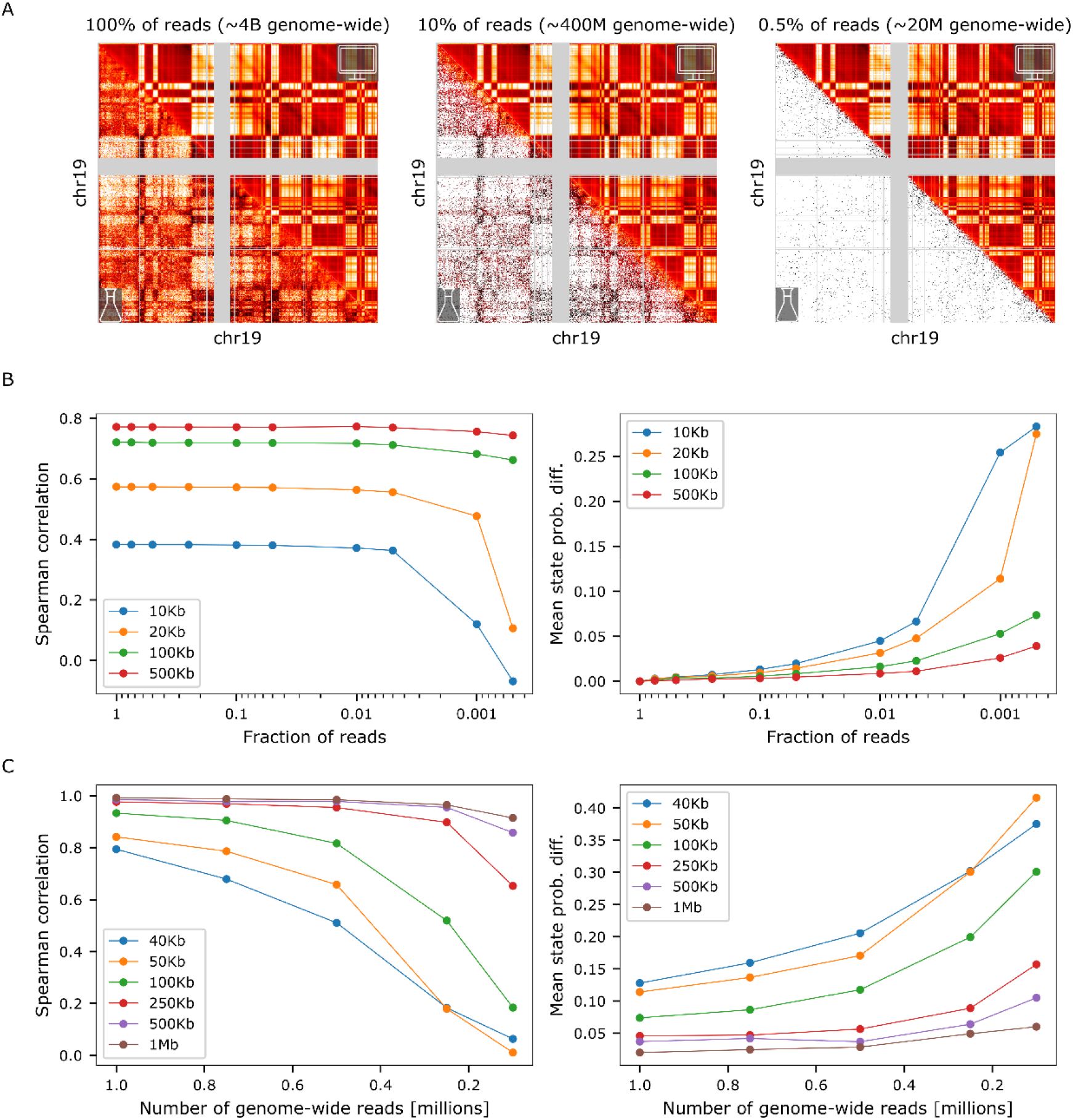
Model robustness at low sequencing depth. (A) Comparison of chromosome 19 interaction frequencies and model inferred interaction probabilities at 20kb resolution when using 100%, 10% and 0.5% of the data. Matrices were distance-normalized. (B) Model performance and stability when fitted to down-sampled chromosome 19 data at various resolutions. Left: Distance-normalized Spearman correlation between the sampled Hi-C interaction frequencies and the model-inferred interaction probabilities. Right: mean absolute difference in state probabilities between a model fitted on the entire data and a model fitted on down-sampled data. (C) Model performance and stability at single-cell Hi-C sequencing depths. We inferred interaction probabilities on chromosome 19 at 20kb resolution, treating these as “true” interaction probabilities, and sampled interactions from these probabilities. Left: Distance-normalized Spearman correlation between the “true” interaction probabilities and the interaction probabilities inferred from the sampled interactions. Right: mean absolute difference between the “true” state probabilities and the state probabilities inferred from the sampled interactions.

Given the ability of the model to perform well on very poorly sampled interaction maps, we asked how well the model would work on single-cell Hi-C maps. To evaluate this, we took the interaction probability matrix predicted from GM12878 chromosome 19, and treated this map as the “true” interaction probabilities. We then randomly sampled interactions from these probabilities, at coverage levels similar to that of single-cell maps (100K-1M interactions genome-wide). Finally, we applied our model to these sampled matrices, allowing us to evaluate the model’s ability to recover the “true” interaction probabilities and state probabilities. Applying our model, we find that the model recovers the “true” parameters (correlation>0.9, mean state probability error<0.1) with as few as 100K reads at 0.5Mb resolution, 250K reads at 0.25Mb resolution, and 750K reads at 0.1Mb resolution (**Figure 3C**). We conclude that our model is applicable to single-cell Hi-C data, provided they are sufficiently sampled.

### Differences in cis and trans interactions of genomic compartments

Trans interactions are less frequent than cis interactions, and in the context of genomic compartments trans interactions are often considered a mere extension of the genomic compartments observed in cis. Indeed, it is common to ignore either trans interactions or cis interactions when analyzing genomic compartments, under the implicit assumption that they follow similar principles. We decided to use our model to revisit analyses performed by Imakaev et al.(54) and test this hypothesis directly. First, we modified our model to cope with whole-genome modelling. Briefly, this involved optimization of the inference method (the whole-genome interaction frequency matrix is more than 150 times larger than the interaction frequency map of the largest chromosome), as well as modifying the distant-dependent interaction term so it still uses a power-law decay in cis but uses a constant background interaction level *β* in trans. After fitting the two-state model to the genome-wide GM12878 interaction map at 100Kb resolution, we find self-affinities of 0.51 for state 1, 0.48 for state 2, and 0.01 affinity between the states (**Figure 4A**). Next, we created a saddle plot of the GM12878 interaction map on chromosomes 1 and 2 by sorting both the rows and columns according to their inferred probability of being in state 1 (**Figure 4B**). We expected the quadrants of the saddle plot to roughly reflect the state affinity matrix, and this is indeed what we observed. Next, we created separate saddle plots for chromosome 1, chromosome 2, and their trans interactions (**Figure 4C**). We observe that the two cis saddle plots are similar to each other with approximately equal self-affinities for state 1 and state 2, while the trans saddle plot is different with state 1 showing notably higher self-affinity than state 2, suggesting that state interaction affinities may differ between cis and trans. To allow our model to account for these differences, we extended our model to use two separate state affinity matrices for cis and trans. Refitting this extended model, we infer cis and trans affinity matrices and find that they match the observed cis and trans saddle plots (**Figure 4D, 4E**), so that state 1 self-affinity in cis is similar to that of state 2 (0.49 state 1 vs. 0.51 state 2) but is much higher in trans (0.66 state 1 vs. 0.27 state 2). Comparing the inferred state probabilities to histone modification data, we find that state 1 is correlated with active chromatin marks (“compartment A”) while state 2 is anti-correlated (“compartment B”). We find qualitatively similar cis and trans affinity matrices when repeating the analysis in additional cell lines (H1-ESC, HFF and mouse ESC). Thus, in line with the observations of Imakaev et al.(54), our results suggest that a whole-genome model of genomic compartments must incorporate different interaction rules in cis and trans, with active chromatin self-associating more frequently in trans than inactive chromatin. The difference between the two states may also allow to distinguish the two states without referring to external datasets as is done in PCA-based compartment analysis.

**Figure 4.**
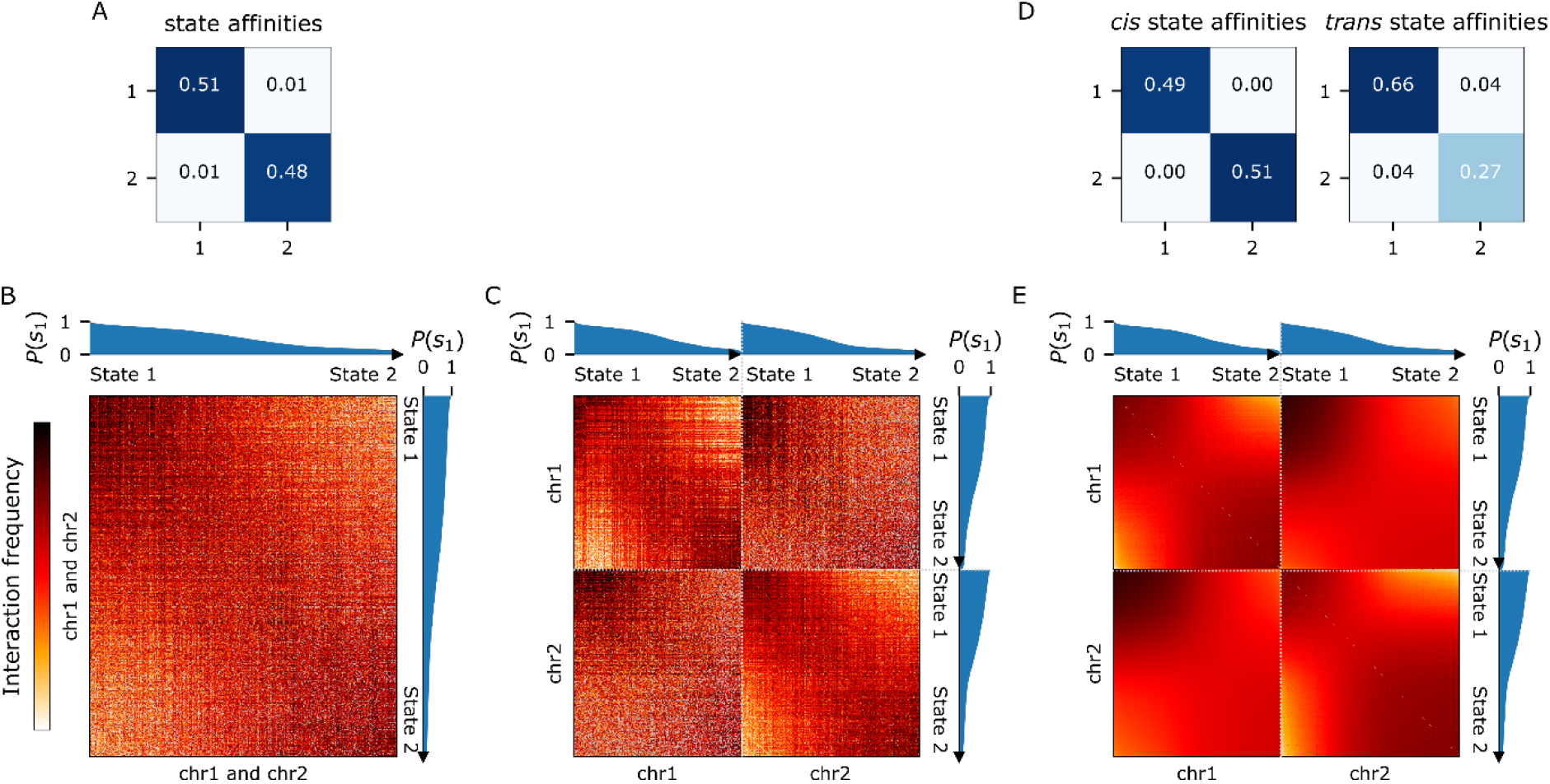
States interact differently within and between chromosomes. (A) State-state affinity matrix from whole genome fit of two-state model at 100Kb resolution. (B) Joint saddle plot of chromosomes 1 and 2. The rows and columns of the Hi-C interaction frequency matrix were sorted by, the probability to be in state 1. Cis and trans data were normalized to have the same mean interaction frequency. (C) Separate saddle plots of chromosomes 1 and 2 in cis and trans. Cis and trans data were normalized to have the same mean interaction frequency. (D) State-state cis and trans affinity matrices from whole genome fit of two-state model with separate affinity matrices (E) Separate saddle plots of chromosomes 1 and 2 in cis and trans using the inferred interaction probabilities rather than the Hi-C interaction frequencies.

### Extending the number of states

We next used the model to investigate the number of states in a principled manner. While a two-state model explained much of the relevant variation in the data, previous work has suggested the possibility of additional compartments or subcompartments(45). We first used our model to identify regions in which a two-state model is clearly insufficient. For example, the chromosomal region shown in **Figure 5A** clearly deviates from a checkered pattern and thus a simple two-state model would not be sufficient, as verified by fitting our two-state model. In order to select a reasonable number of states, we first fitted the entire genome using 2-8 states at 50kb resolution. Next, we calculated for each model the mean reconstruction error for the interaction profile of every genomic bin (a row/column in the interaction map). We reasoned that if the number of states is too low, there would be some subset of rows that would show a high reconstruction error. We thus examined the standard deviation of the mean row reconstruction error as a function of the number of states in the model, expecting the standard deviation to decrease as the number of states increases. The resulting plot exhibited knee points at four and seven states (**Figure 5B**). Preferring a simpler model, we chose to proceed with a four-state model. Revisiting the example region in **Figure 5A**, we find that the four-state model is far superior in capturing the observed pattern. Finally, comparing the two-state and four-state genome-wide models (**Figure 5C**), we find that the four-state model achieves a higher Spearman correlation (two-state: 0.64, 0.1 s.d.; four-state: 0.64, 0.1 s.d.) and is closer to the optimal possible correlation (two-state: 0.15, 0.08 s.d.; four-state: 0.11, 0.04 s.d.). Our results suggest that at least four states may be needed in order to adequately describe genomic compartments.

**Figure 5.**
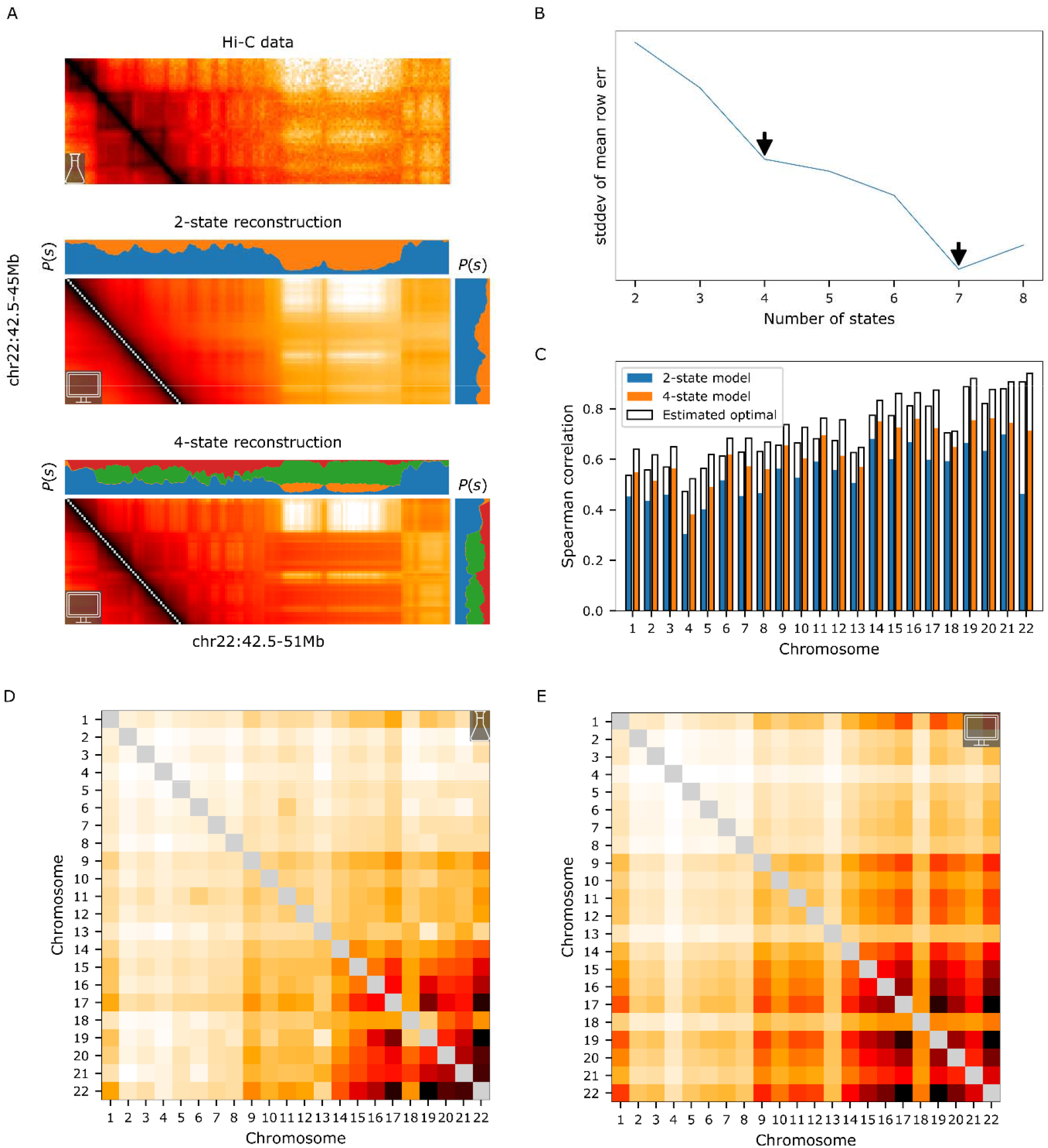
Extending the model beyond two states. (A) Modelling a complex region in chromosome 22 (50Kb resolution). Top: Hi-C interaction frequencies. Middle: interaction probabilities inferred by a two-state model, accompanied by locus state probabilities (blue state 1, orange state 2). Bottom: Middle: interaction probabilities inferred by a two-state model, accompanied by locus state probabilities (blue state 1, orange state 2, green state 3, red state 4) (B) Standard deviation of the mean row reconstruction error as a function of the number of states in the model. Arrows indicate knee points at four and seven states. (C) Distance-normalized Spearman correlation between the Hi-C interaction frequency matrix and the model’s inferred interaction probability matrix, for whole-genome two-state and four-state models. The optimal possible correlation for each model at matching resolution and sequencing depth is shown as reference (see Methods for details). (D) GM12878 chromosome-level interaction frequency matrix. (E) chromosome-level interaction probability matrix inferred by whole-genome four-state model.

It has been shown that on the whole chromosome level, while the relative nuclear positions of chromosomal territories are highly stochastic, certain pairs of chromosomes tend to interact more frequently with others (1, 78). Polymer simulations have suggested that chromatin activity-based segregation may be sufficient to explain chromosome nuclear positioning (79). We asked whether chromosome-level interactions can be explained by our model or whether other mechanisms are involved. We thus calculated a chromosome-level interaction map for GM12878, reobserving the known tendency of small gene-rich chromosome to interact. We then used our 4-state model to predict a chromosome-level interaction probability map. We find that the predicted and observed matrices correspond well (Spearman correlation 0.93) (**Figure 5D, 5E**), suggesting that chromosome-level interaction can be largely explained by local state-based interaction.

### Interpreting model parameters

Following the selection of the four-state model, we turned to inspect the properties of the inferred parameters including the state affinity matrices and the locus state probabilities. Interestingly, we find that the cis state affinity matrix is diagonal, suggesting little interaction between any of the states (**Figure 6A**). In contrast to the two-state model, we observe differences in state self-affinities (0.16, 0.17, 0.32, 0.35 for states 1, 2, 3, 4 respectively). However, we find that the trans state affinity matrix is notably different from that of cis (**Figure 6B**): The matrix is no longer diagonal, albeit off-diagonal values are low and could be attributed to noise (<=0.06), and the state self-affinities also differ considerably, e.g. state 1 has the lowest self-affinity in cis (0.16) but the highest self-affinity in trans (0.32). Examining locus state probabilities, we find that each state displays a distinct multimodal distribution of probabilities, suggesting that the states are not redundant (**Figure 6D**). This is further supported by calculating the correlations between all pairs of states, demonstrating no pair of states is highly correlated (**Figure 6E**). Taken together, these results support the notion of four individual states, each with distinctive cis and trans affinities governing interactions.

**Figure 6.**
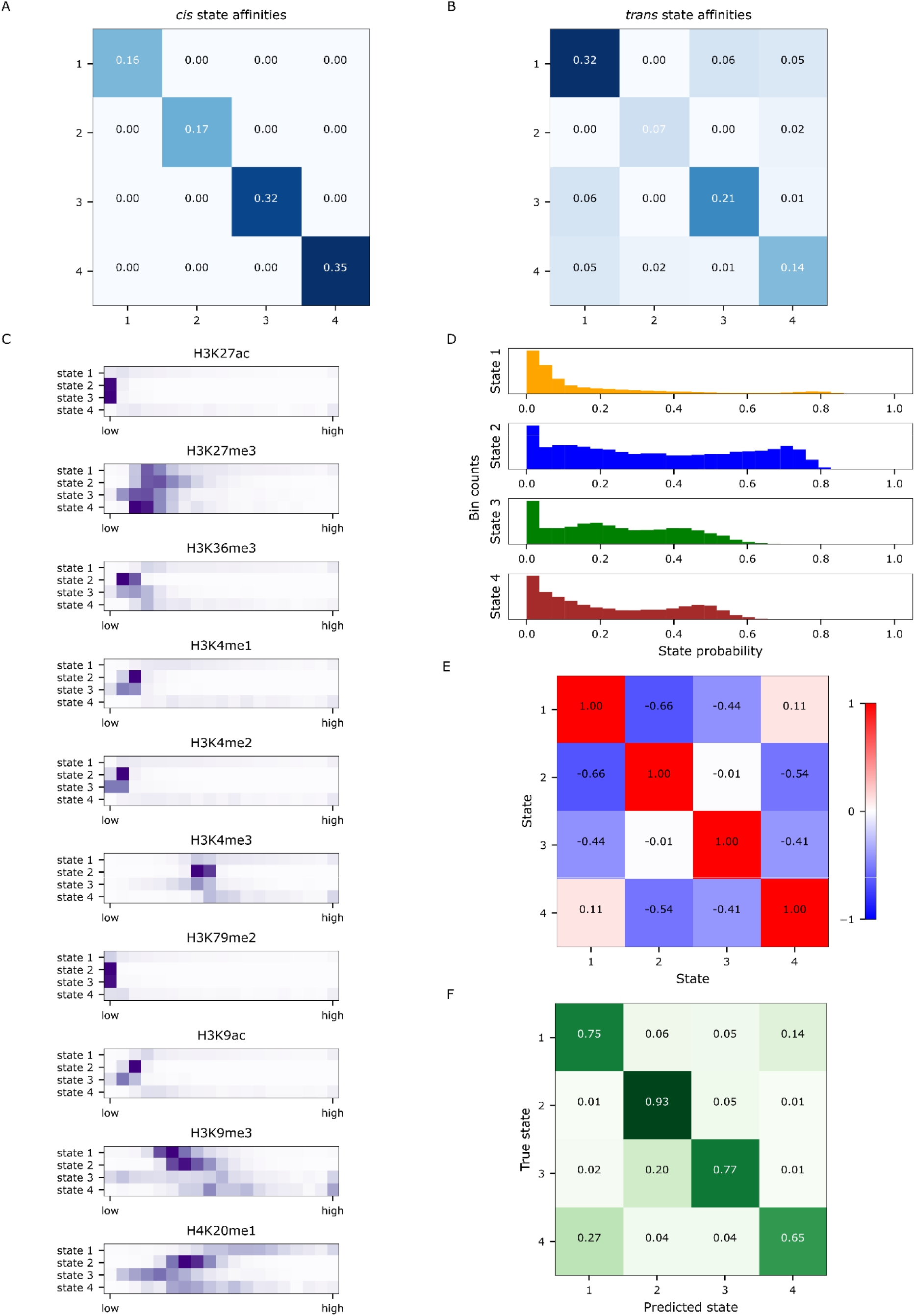
Analysis of four-state model parameters. All results shown were taken from fitting the whole-genome four-state model at 50Kb resolution. (A) State affinity cis matrix. (B) State affinity trans matrix. affinities matrices for the 4-state fit at 50Kb resolution showing different *cis* and *trans* affinities for all states. (C) Heatmaps representing the distribution of histone modification frequency for ten different ENCODE (80) ChIP-Seq histone modification tracks, separated by state. (D) Histograms of locus state probabilities genome-wide. (E) Pearson correlation matrix of locus state probabilities. (F) Confusion matrix depicting locus state prediction by an elastic net multinomial logistic regression classifier from locus histone modifications

### Histone modifications underlie interaction states

We next asked what molecular markers may underlie the four states inferred by our model. We assembled ten ENCODE (80) GM12878 histone modification ChIP-Seq tracks. As our model assigns a state probability to each genomic locus (50kb bin), we first assembled for each state the set of bins in which that state has a probability of at least 0.6 This yielded 4667, 13136, 694 and 327 bins for states 1, 2, 3 and 4 respectively. We first examined visually the distributions of each of the histone modifications for each of the four states (**Figure 6C**). For each of the histone modifications, we see potentially informative differences between the states, with the most prominent differences in the distributions of H4K20me1. In spite of these aggregate differences, it is unclear to what extent these could be used to predict locus state based on locus histone modifications. We thus addressed this question directly by training an elastic net multinomial logistic regression classifier to predict locus state from histone modifications. Training the classifier on odd-numbered chromosomes and testing on even-numbered, we obtain an overall test accuracy of 0.87, with classification errors tending to occasionally misclassify state 3 as 2 and state 4 as 1 (**Figure 6F**). We thus conclude that the four inferred states are indeed distinct and are marked with characteristic combinations of histone modifications.

### Single-cell analysis of state mixing

Lastly, we asked whether the mixing of states at a given locus, which is an inherent feature of our model, occurs at the population or cell level. For example, consider a locus which is inferred to be 50% state 1 and 50% state 2. In population-level mixing, the apparent mixing of states results from the equal mixing of two populations, one in which the locus is completely in state 1 and the other in which the locus is completely in state 2. An alternative scenario is cell-level mixing, consisting of one homogenous population in which the locus itself is in a mixed intermediate state at the single-cell level. While we derived the model with population-level mixing in mind, the formulation of our model cannot distinguish between these two scenarios. To address this question, we identified within the GM12878 interaction map an area in chromosome 19 which shows evidence of state mixing (**Figure 7A**). Within this area, we observed a genomic region whose interaction pattern appears to be a mixing of the interaction patterns of two nearby regions. We first checked that our model identifies this pattern as mixed, and found that the seven-state model indeed identifies one region as higher state 3, another as higher state 7, and the mixed region as a mix of 3+7 (**Figure 7A**). Next, we elected to use 1129 single-cell Hi-C maps measured by Kim et al. (81) to distinguish between population-level and cell-level mixing. We reasoned that if the mixing of the 3+7 region is cell-level, a single-cell interaction profile of the 3+7 region would be more likely to resemble the bulk interaction profile of the mixed 3+7 region than the bulk interaction profile of state 3 or state 7 regions (**Figure 7B, 7C**). If the mixing of the 3+7 region is population level, the opposite would be more likely. We verified the plausibility of this approach by simulating population and cell-level mixing scenarios (see Methods). Single-cell interaction profiles were sampled either from the bulk mixed 3+7 region interaction profile (cell-level mixing), or from the two bulk 3/7 state region interaction profiles (population-level mixing, 50% each state). We then calculated for each simulated single-cell interaction profile the log ratio between its likelihood of coming from the mixed state and its likelihood of coming from one of the pure states. This log ratio was further normalized to account for expected differences in distance-dependent interaction between the genomic regions (see Methods), and is hereon referred to as Mixing Log Ratio (MLR). Examining the resulting MLR distributions in the two simulated mixing scenario (**Figure 7D**), we find that the MLRs in the cell-level mixing simulation tend to be positive (median 0.76) while the MLRs in the population-level mixing simulation tend to be negative (median -0.96), suggesting the two scenarios can be distinguished with this approach (one-sided Kolmogorov-Smirnov p-value<10^−193^). Finally, to examine which of the two scenarios better fits the data, we calculated the MLRs for the measured single cell Hi-C data. We find that the resulting MLRs tend to be negative (median -1.1) and the distribution overall is more consistent with the population-level mixing scenario (two-sided Kolmogorov-Smirnov p-value<10^−160^). While not excluding the possibility of cell-level mixing occurring elsewhere, our results suggest that the observed mixed pattern is mostly due to population-level mixing.

**Figure 7.**
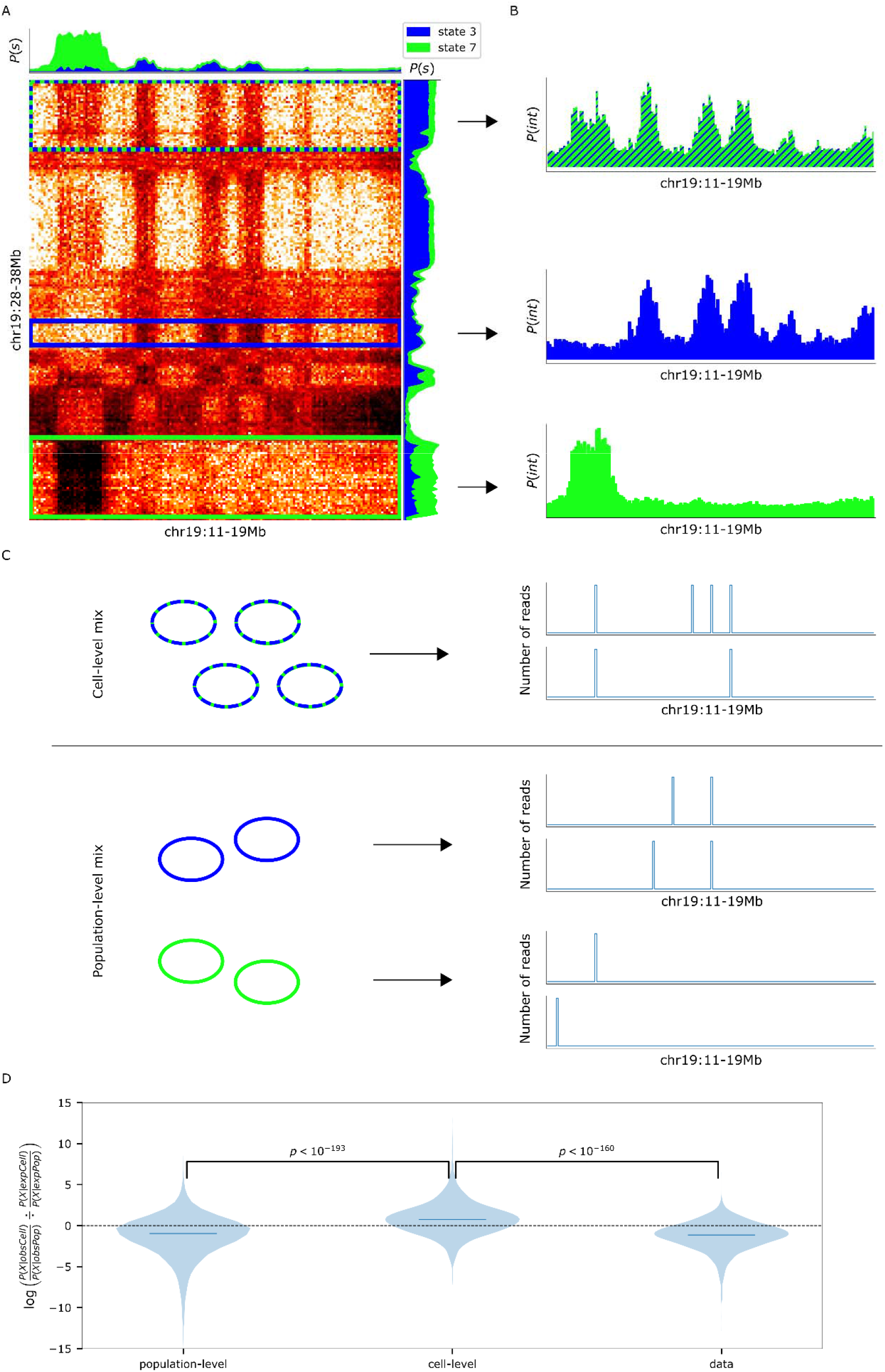
Analysis of state mixing. (A) Evidence of state mixing in chromosome 19. The interaction pattern marked in the green-blue rectangle appears to be a mix of the interaction pattern marked in the blue rectangle and the in the green rectangle. Locus state probabilities *P*(*s*) are shown for states 3 and 7 taken from the seven-state model. (B) Averaged interaction profiles for the green-blue, blue, and green regions. (C) Schematic of simulated single-cell Hi-C profiles generated by the two simulated mixing scenarios. Top: In population-level mixing, sparse single-cell interaction profiles are sampled from the previously shown green-blue interaction profile. Bottom: In cell-level mixing, sparse single-cell interaction profiles are sampled 50% from the previously shown green interaction profile and 50% from the blue interaction profile. (D) Violin plots of the distributions of the Mixing Log Ratio for cell-level mixing simulation, population-level mixing simulation, and real single-cell Hi-C data from Kim et al. (81). Mixing Log Ratio represents the logarithm of the ratio between the likelihood of a single-cell profile given cell-level mixing and the likelihood of the single-cell profile given population-level mixing, after accounting for expected distance-dependent differences (see Methods). Kolmogorov-Smirnov p-values are shown.

## Discussion

In this work, we present deGeco, a probabilistic modelling approach rooted both in data-driven and hypothesis-driven approaches. On one hand, this provides us with an explicit model that has biologically interpretable parameters and can be used to test biological hypotheses by simple modifications. On the other hand, this enables us to utilize the power of the large amounts of available data to solve inverse-type problems, such as estimating the locus states across the genome or the affinities between different states within and between chromosomes.

We evaluate the robustness of our model with respect to the amount of sequencing reads, as sequencing depth of Hi-C libraries is still cost-prohibitive due to the huge space of possible pairwise interactions. We show that approximately 20M read pairs are sufficient to accurately infer model parameter values which are very close to those inferred from a 4000M read pair map, even at 10Kb bin resolution. It is interesting to consider this in the context of several recent methods for reconstructing interaction maps from very sparse data by first training deep-learning models on well-sampled high-resolution maps (74–77). Although our method is currently limited to genomic compartments and larger interaction patterns, it is notable that simply by the virtue of its few mechanistic assumptions, it is able to perform this type of reconstruction from very little data without training first on any data. We also try to push the limits of the method by attempting to apply it to single-cell Hi-C maps, and find it can correctly infer genomic compartments at lower resolutions (250-500kb) for reasonably well-sampled single-cell maps.

Due to the size of mammalian Hi-C interaction matrices, which poses a significant computational hurdle, computational methods often resort to operating on smaller scales, such as single chromosome interactions or small genomic windows. This often carries the implicit assumption that the rules for interaction are consistent across the genome. We enabled whole-genome inference with our model by developing an optimized sparse representation coupled with a sampling-based estimation of the partition function (see Methods). We then used this to interrogate the entire genome simultaneously, and surprisingly found that state-state affinities between chromosomes differ considerably from those within chromosomes. In addition to the implications of these findings for future genome-wide models, it would be interesting to further explore the physical basis of these differences by using complementary modelling techniques, especially those based on polymer models.

Since “all models are wrong, but some are useful”, it is often useful to inspect where a model was wrong. Although a two-state model explained most of the explainable variance in the GM12878 interaction map, we used the model to identify regions which were clearly not explained well by two states. This led us to evaluate the number of states, finding that four and seven states might be reasonable choices. Naturally, other choices are possible, and these could change between cell types and species. Although several others have proposed that more than two states should be considered (9, 45, 55, 56, 62, 66, 69), without an explicit model it is difficult to distinguish an interaction state from an interaction pattern. For example, a mixing of states could create an interaction pattern that would seem different from those of the individual states, and would be falsely identified as a separate state by approaches such as clustering. Pursuing the molecular basis of the four-state model, we find characteristic state probabilities and histone marks for each of these, ultimately constructing a simple classifier that predicts locus state from histone marks with 87% accuracy. Although it has been shown that chromatin features can be separated into states(82, 83), and that these can be used to predict spatial interactions(9, 67), it remains to be seen whether there is a simple mapping between chromatin states and interaction states.

Finally, we decided to investigate one of the assumptions of the model, namely that within single cell every locus is in exactly one state so that any mixing of states at a locus is due to population-level mixing (averaging over a heterogenous cell population). However, if we would have changed the assumption so that even within a single cell a locus can be in multiple states, e.g. interacting simultaneously like two other states, the model formulation would be the same. We thus decided to use single-cell Hi-C data to find whether there is evidence for cell-level mixing, and coupled with probabilistic simulations conclude that we do not find evidence of cell-level mixing. In this respect, it is important to note that cell-level mixing can appear artificially due to genomic bin size: if loci of different states occupy the same genomic bin, the bin might seem to interact simultaneously with multiple states.

In conclusion, we envision this approach as a probabilistic modelling framework for further hypothesis-driven investigation and interpretation of genomic compartments as well as other interaction patterns such as TADs and point interactions.

## Materials and methods

### Full model description

Based on the assumptions described in the Results section, we propose the following probabilistic model for pairwise interaction probability between loci:

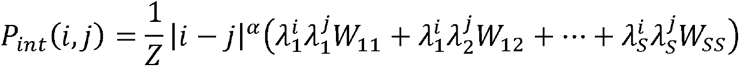

Where 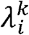 is the probability over the cell population that locus *i* is in the *k*-th state (we assume 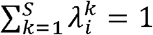) *W*_*km*_ is a non-negative affinity between states *k* and *m* (we assume *W*_*km*_ = *W*_*mk*_ and ∑_*k,m*,_ *W*_*km*_ = 1); |*i* − *j* |^*α*^ is a distance-dependent interaction function (we assume *α* < 0); and *Z* is a normalization factor which ensures ∑_*i,j*_ *P*_*int*_ (*i, j*) = 1, also known as the partition function. Locus interactions are symmetric, so *P*_*int*_(*i,j*) = *P*_*int*_(*i,j*). We note that if we define a matrix Λ of locus state probabilities such that 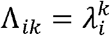 the genomic compartments component can be rewritten as matrix factorization:

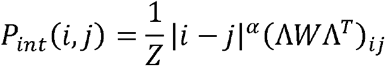

For the genome-wide model (see Results), we modify the distance-dependent function interactions such that if *i,j* are in cis *d* = |*i* − *j* |^*α*^ and if they are in trans *d* = *β* (*β* is a parameter representing the general strength of trans interactions). As described in the Results section, we later used separate *W* matrices for cis and trans.

Since we do not have access to the true interaction probabilities, we use Hi-C interaction frequencies to estimate them. We assume that *R* Hi-C read pairs are multinomially sampled from the interaction probability distribution *P*_*int*_, yielding the interaction frequency matrix *X* (so ∑_*i*≤*j*_ *X*_*ij*_ = *R*). Thus, the probability of obtaining Hi-C interaction frequency matrix *X* is:

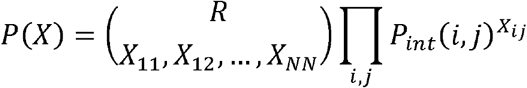

Where *N* is the number of bins in the matrix and *P*_*int*_ (*i, j*) is specified by the model. The log-likelihood of the full model is then given by:

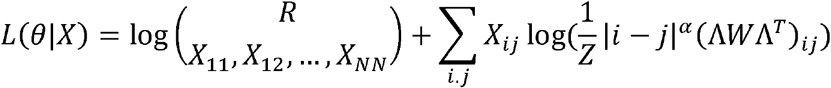

Where *θ* are the model parameters *α*, Λ and *W*. In order to estimate the values of the parameters given a Hi-C interaction matrix *X*, we maximize this log-likelihood objective function. Note that given *X* the multinomial coefficient is constant and can be ignored during maximization.

### Optimizing the objective function

Unfortunately, the objective function is not concave, requiring multiple initializations. To direct the search towards good solutions, we developed a multi-resolution fitting strategy consisting of running the model at a low resolution (large bin size) multiple times, and proceeding with the best solutions as initialization points for refinement at higher resolutions (small bin sizes). The underlying logic is that a good solution at high resolution must be a good solution in low resolution. Note that when moving to a higher resolution, the locus state probabilities are duplicated according to the ratio of resolutions, for example the state probability inferred for a 500Kb bin is duplicated into five 100Kb bins. Unless stated otherwise, the model was run 20 times at 500Kb with random seeds 1-20, the 5 best solutions (by likelihood) were refined at 100Kb and then at 50Kb. We use the best solution (by likelihood) out of the 5 refined solutions.

Since the Hi-C matrix is very large in high resolutions, especially when considering a whole genome, the memory and CPU requirements quickly become prohibitive. To address this, we developed a strategy for leveraging sparse data structures and partition function estimation. Our strategy is based on two observations. First, Hi-C interaction frequency matrices, especially in high resolution, are sparse as they are mostly populated with zeros. Second, when looking at the sum ∑_*i,j*_ *X*_*ij*_ log *P*_*int*_(*i, j*) in the objective function, zero entries in *X* cancel out most of the sum terms and obviates computation of the corresponding entries in *P*_*int*_. However, *P*_*int*_(*i, j*) contains the normalization term (partition function) *Z* = ∑*P*_*int*_(*i, j*), which requires information from all entries, even those with corresponding zero values in *X*. To work around this, we note that *Z* can be viewed as a sum of two groups of entries, those with corresponding non-zero values in *X* and those with corresponding zero values: *Z* = ∑_*nonzeros*_ *P*_*int*_(*i, j*) ∑_*zeros*_ *P*_*int*_(*i, j*). We calculate ∑_*nonzeros*_ *P*_*int*_(*i, j*) exactly, and estimate ∑_*zeros*_ *P*_*int*_(*i, j*) by taking a random sample of entries with corresponding zero values in *X* (without replacement). Thus, our implementation dramatically reduces memory and CPU requirements by using the Hi-C matrix in sparse form where only non-zero entries are held in memory, and calculating *P*_*int*_(*i, j*) only for entries corresponding with non-zero *X* entries or corresponding with a random subset zero of *X* zero entries. We chose to use a number of sampled non-zero entries equivalent to the number of zero entries.

### Datasets

Human genome hg38 version was used. GM12878 Hi-C matrix by Rao et al.(45) was obtained from the 4D Nucleome data portal (84) and used in all analyses unless specified otherwise. The matrix was balanced and binned by cooler (85). Analyses were not performed on chromosomes X and Y. GM12878 single-cell Hi-C data by Kim et al. (81) was obtained from https://noble.gs.washington.edu/proj/schic-topic-model/. Histone modification GM12878 ChIP-Seq data tracks (fold-change over input) were obtained from ENCODE (80) for H3K27ac, H3K27me3, H3K36me3, H3K4me1, H3K4me2, H3K4me3, H3K79me2, H3K9ac, H3K9me3 and H4K20me1. Histone tracks were binned to 50Kb bins by taking the average fold-change over input across the entire bin, considering missing values as zero. To reduce effects of outliers, every histone track had its highest 1% of values trimmed to the 99^th^ percentile level.

### Model robustness analysis

For the analysis in Figure 3B: Interactions were sampled from the unbalanced GM12878 Hi-C matrix of Chromosome 19 without replacement, followed by balancing and binning (where relevant) by cooler (85). To fit the model at various resolutions, the model was run with 10 random initializations at 500Kb resolution, and these were further refined to 100Kb, 20Kb and 10Kb resolutions. To evaluate performance, the best solution (by likelihood) was chosen for each resolution.

For the analysis in Figure 3C: Interaction probabilities and locus state probabilities were inferred by a two-state model fitted to GM12878 Hi-C matrix of chromosome 19. These were treated as the “true” probabilities, and we tested the ability of the model to recover these from extremely sparse data. Sparse interaction maps were generated by multinomially sampling from the “true” interaction probabilities. Sampling level matched 0.1-1Mb genome-wide interactions, similar to the sequencing depth achieved in recent single-cell Hi-C maps. To fit the model at various resolutions, the model was run with 10 random initializations at 1Mb resolution, and these were further refined to consecutively higher resolutions. To evaluate performance, the best solution (by likelihood) was chosen for each resolution.

### Analysis of histone modifications

Analysis of histone modifications was performed at 50Kb resolution. To avoid complications due to state mixing, analyses were performed only on bins in which one state is dominant (state probability>0.6). This resulted in 4667, 13136, 694 and 327 bins for states 1, 2, 3 and 4. To classify the locus state for each of these bins from the locus histone modifications data, chromosomes were split into test and train sets, which contained the odd and even chromosomes, respectively. The training set was scaled to zero mean and unit standard deviation, and the test set was scaled according to the mean and standard deviation learned from the training set. To avoid effects due to classes 3 and 4 being much smaller, their samples in the training set were each duplicated 15 times. No duplication was done on the test set. A multiclass logistic regression classifier was trained using the multinomial loss and elastic net regularization (scikit-learn(86) implementation). The C (inverse of regularization strength) and L1/L2 ratio hyperparameters were optimized using 10-fold cross validation on the training set and searching over a 10×10 grid in the ranges (0,1) for C and (1e-4,1e4) for L1/L2 ratio. The overall accuracy was calculated as the fraction of correct classifications in the test set. To offset differences in the sizes of the classes in the test set, the confusion matrix was row-normalized.

### State mixing analysis

To simulate state-mixing scenarios in single cells, we first created average interaction probability profiles for three groups shown in Figure 7A: a region with higher state 3 (blue), a region with higher state 7 (green) and a mixed 3+7 region (green-blue). To this end, we collected rows in the Hi-C matrix with a similar interaction profile to each of the three regions from the entire chromosome. To reduce the effects of genomic distance, we considered only interactions at >2Mb distance in all mixing analyses. Finally, the rows of each of the three groups were averaged and converted to probabilities by normalizing to one, adding 1e-10 as pseudocounts, and renormalizing, producing probability vectors *p*^3^, *p*^7^ and *p*^3+7^. In addition, we created probability vectors for each of the three groups based only on distance-dependent interaction signal (sometimes referred to as “expected” signal), by replacing every entry in the Hi-C matrix with the average interaction frequency of its diagonal. We refer to these as *p*^*exp*3^, *p*^*exp*7^and *p*^*exp*3+7^.

To simulate single-cell interaction profiles from each of the scenarios, we used the scHi-C experiment GM12878 cells by Kim et al. (81). Single-cell maps were used at 500kb resolution due to their sparsity. Out of 4545 cells, maps of 1129 cells were chosen which had at least five reads in the mixed region in chromosome 19. We started by calculating the average interaction frequency profiles within the mixed region for each of the single-cell maps, yielding 1129 interaction frequency vectors *X*^*data*^. We first simulated cell-level mixing, by sampling 1129 interaction frequency vectors *X*^*cell*^ multinomially from *p*^3+7^, matching the number of sampled to reads in each vector to the number of reads in the respective vector in *X*^*data*^. To simulate population-level mixing, we sampled 1129 interaction frequency vectors *X*^*pop*^, where half are sampled multinomially from *p*^3^ and the other half are sampled from *p*^7^, again matching the number of reads to *X*^*data*^.

To quantify whether a single-cell interaction frequency vector *x* is more likely to come from cell-level or population-level mixing, we first note that the probability of observing an interaction frequency vector *x* given a probability vector *p* is:

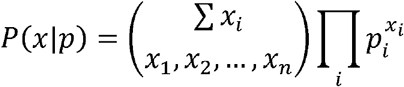

Next, we define the Mixing Log Ratio (MLR) for an interaction frequency vector *x* to be:

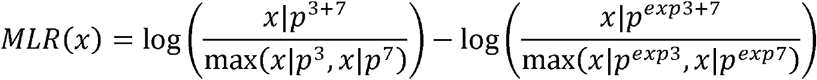

Finally, we calculated MLR value for *X*^*cell*^, *X*^*pop*^ and *X*^*data*^.

## Data availability

Code was written in Python and Cython (87) mainly using the SciPy and NumPy (88) libraries. Code is available at https://github.com/KaplanLab/deGeco.

## Acknowledgements

We thank the members of the Kaplan Lab for helpful comments and discussions. We thank David Cohen for HPCC support and administration.

## References

1. Lieberman-Aiden, E., van Berkum, N.L., Williams, L., Imakaev, M., Ragoczy, T., Telling, A., Amit, I., Lajoie, B.R., Sabo, P.J., Dorschner, M.O., et al. (2009) Comprehensive mapping of long-range interactions reveals folding principles of the human genome. Science, 326, 289–93.

2. Quinodoz, S.A., Ollikainen, N., Tabak, B., Palla, A., Schmidt, J.M., Detmar, E., Lai, M.M., Shishkin, A.A., Bhat, P., Takei, Y., et al. (2018) Higher-Order Inter-chromosomal Hubs Shape 3D Genome Organization in the Nucleus. Cell, 174, 744-757.e24.

3. Mccord, R.P., Kaplan, N. and Giorgetti, L. (2020) 3C and beyond□: towards an integrative view of chromosome structure and function. Mol. Cell, 10.1016/j.molcel.2019.12.021.

4. Zhao, Z., Tavoosidana, G., Sjölinder, M., Göndör, A., Mariano, P., Wang, S., Kanduri, C., Lezcano, M., Sandhu, K.S., Singh, U., et al. (2006) Circular chromosome conformation capture (4C) uncovers extensive networks of epigenetically regulated intra- and interchromosomal interactions. Nat. Genet., 38, 1341–7.

5. Denker, A. and Laat, W. De (2016) The second decade of 3C technologies□: detailed insights into nuclear organization. 10.1101/gad.281964.116.

6. Guelen, L., Pagie, L., Brasset, E., Meuleman, W., Faza, M.B., Talhout, W., Eussen, B.H., De Klein, A., Wessels, L., De Laat, W., et al. (2008) Domain organization of human chromosomes revealed by mapping of nuclear lamina interactions. Nature, 453, 948–951.

7. Beagrie, R.A., Scialdone, A., Schueler, M., Kraemer, D.C.A., Chotalia, M., Xie, S.Q., Barbieri, M., De Santiago, I., Lavitas, L.M., Branco, M.R., et al. (2017) Complex multi-enhancer contacts captured by genome architecture mapping. Nature, 543, 519–524.

8. Lupiáñez, D.G., Kraft, K., Heinrich, V., Krawitz, P., Brancati, F., Klopocki, E., Horn, D., Kayserili, H., Opitz, J.M., Laxova, R., et al. (2015) Disruptions of topological chromatin domains cause pathogenic rewiring of gene-enhancer interactions. Cell, 161, 1012–1025.

9. Rowley, M.J., Nichols, M.H., Lyu, X., Ando-Kuri, M., Rivera, I.S.M., Hermetz, K., Wang, P., Ruan, Y. and Corces, V.G. (2017) Evolutionarily Conserved Principles Predict 3D Chromatin Organization. Mol. Cell, 67, 837-852.e7.

10. Hafner, A. and Boettiger, A. (2022) The spatial organization of transcriptional control. Nat. Rev. Genet., 10.1038/s41576-022-00526-0.

11. Le Dily, F., Baù, D., Pohl, A., Vicent, G.P., Serra, F., Soronellas, D., Castellano, G., Wright, R.H.G., Ballare, C., Filion, G., et al. (2014) Distinct structural transitions of chromatin topological domains correlate with coordinated hormone-induced gene regulation. Genes Dev., 28, 2151–62.

12. Sima, J., Chakraborty, A., Dileep, V., Michalski, M., Klein, K.N., Holcomb, N.P., Turner, J.L., Paulsen, M.T., Rivera-Mulia, J.C., Trevilla-Garcia, C., et al. (2018) Identifying cis Elements for Spatiotemporal Control of Mammalian DNA Replication. Cell, 0, 1–15.

13. Pope, B.D., Ryba, T., Dileep, V., Yue, F., Wu, W., Denas, O., Vera, D.L., Wang, Y., Hansen, R.S., Canfield, T.K., et al. (2014) Topologically associating domains are stable units of replication-timing regulation. Nature, 515, 402–405.

14. Yaffe, E., Farkash-amar, S., Polten, A., Yakhini, Z., Tanay, A. and Simon, I. (2010) Comparative Analysis of DNA Replication Timing Reveals Conserved Large-Scale Chromosomal Architecture. PLoS Genet., 6.

15. Giorgetti, L., Lajoie, B.R., Carter, A.C., Attia, M., Zhan, Y., Xu, J., Chen, C.J., Kaplan, N., Chang, H.Y., Heard, E., et al. (2016) Structural organization of the inactive X chromosome in the mouse. Nature, 535, 575–9.

16. Nora, E.P., Lajoie, B.R., Schulz, E.G., Giorgetti, L., Okamoto, I., Servant, N., Piolot, T., van Berkum, N.L., Meisig, J., Sedat, J., et al. (2012) Spatial partitioning of the regulatory landscape of the X-inactivation centre. Nature, 485, 381–385.

17. Crane, E., Bian, Q., McCord, R.P., Lajoie, B.R., Wheeler, B.S., Ralston, E.J., Uzawa, S., Dekker, J. and Meyer, B.J. (2015) Condensin-driven remodelling of X chromosome topology during dosage compensation. Nature, 523, 240–244.

18. Deng, X., Ma, W., Ramani, V., Hill, A., Yang, F., Ay, F., Berletch, J.B., Blau, C.A., Shendure, J., Duan, Z., et al. (2015) Bipartite structure of the inactive mouse X chromosome. Genome Biol., 16, 152.

19. Bonev, B., Mendelson Cohen, N., Szabo, Q., Fritsch, L., Papadopoulos, G.L., Lubling, Y., Xu, X., Lv, X., Hugnot, J.P., Tanay, A., et al. (2017) Multiscale 3D Genome Rewiring during Mouse Neural Development. Cell, 171, 557-572.e24.

20. Ke, Y., Xu, Y., Chen, X., Feng, S., Liu, Z., Sun, Y., Yao, X., Li, F., Zhu, W., Gao, L., et al. (2017) 3D Chromatin Structures of Mature Gametes and Structural Reprogramming during Mammalian Embryogenesis. Cell, 170, 367-381.e20.

21. Phillips-Cremins, J.E., Sauria, M.E.G., Sanyal, A., Gerasimova, T.I., Lajoie, B.R., Bell, J.S.K., Ong, C.-T., Hookway, T.A., Guo, C., Sun, Y., et al. (2013) Architectural protein subclasses shape 3D organization of genomes during lineage commitment. Cell, 153, 1281–95.

22. Stadhouders, R., Filion, G.J. and Graf, T. (2019) Transcription factors and 3D genome conformation in cell-fate decisions. Nature, 569, 345–354.

23. Gibcus, J.H., Samejima, K., Goloborodko, A., Samejima, I., Naumova, N., Nuebler, J., Kanemaki, M.T., Xie, L., Paulson, J.R., Earnshaw, W.C., et al. (2018) A pathway for mitotic chromosome formation. Science (80-.)., 359, eaao6135.

24. Naumova, N., Imakaev, M., Fudenberg, G., Zhan, Y., Lajoie, B.R., Mirny, L.A. and Dekker, J. (2013) Organization of the Mitotic Chromosome. Science, 342, 948–53.

25. Zhang, H., Emerson, D.J., Gilgenast, T.G., Titus, K.R., Lan, Y., Huang, P., Zhang, D., Wang, H., Keller, C.A., Giardine, B., et al. (2019) Chromatin structure dynamics during the mitosis-to-G1 phase transition. Nature, 576, 158–162.

26. Oomen, M.E., Hedger, A.K., Watts, J.K. and Dekker, J. (2020) Detecting chromatin interactions between and along sister chromatids with SisterC. Nat. Methods, 17, 1002–1009.

27. Mitter, M., Gasser, C., Takacs, Z., Langer, C.C.H., Tang, W., Jessberger, G., Beales, C.T., Neuner, E., Ameres, S.L., Peters, J.M., et al. (2020) Conformation of sister chromatids in the replicated human genome. Nature, 586, 139–144.

28. Wang, Y., Wang, H., Zhang, Y., Du, Z., Si, W., Fan, S., Qin, D., Wang, M., Duan, Y., Li, L., et al. (2019) Reprogramming of Meiotic Chromatin Architecture during Spermatogenesis. Mol. Cell, 73, 547-561.e6.

29. Alavattam, K.G., Maezawa, S., Sakashita, A., Khoury, H., Barski, A., Kaplan, N. and Namekawa, S.H. (2019) Attenuated chromatin compartmentalization in meiosis and its maturation in sperm development. Nat. Struct. Mol. Biol., 26, 175–184.

30. Patel, L., Kang, R., Rosenberg, S.C., Qiu, Y., Raviram, R., Chee, S., Hu, R., Ren, B., Cole, F. and Corbett, K.D. (2019) Dynamic reorganization of the genome shapes the recombination landscape in meiotic prophase. Nat. Struct. Mol. Biol., 26, 164–174.

31. Vara, C., Paytuví-Gallart, A., Cuartero, Y., Le Dily, F., Garcia, F., Salvà-Castro, J., Gómez-H, L., Julià, E., Moutinho, C., Aiese Cigliano, R., et al. (2019) Three-Dimensional Genomic Structure and Cohesin Occupancy Correlate with Transcriptional Activity during Spermatogenesis. Cell Rep., 28, 352-367.e9.

32. Belton, J.-M., McCord, R.P., Gibcus, J.H., Naumova, N., Zhan, Y. and Dekker, J. (2012) Hi-C: a comprehensive technique to capture the conformation of genomes. Methods, 58, 268–76.

33. Lajoie, B.R., Dekker, J. and Kaplan, N. (2014) The Hitchhiker’s Guide to Hi-C Analysis: Practical guidelines. Methods, 72, 65–75.

34. Cremer, T. and Cremer, M. (2010) Chromosome territories. Cold Spring Harb. Perspect. Biol., 2, a003889.

35. Lajoie, B.R., Dekker, J. and Kaplan, N. (2015) The Hitchhiker’s guide to Hi-C analysis: practical guidelines. Methods, 72, 65–75.

36. Mirny, L. a. (2011) The fractal globule as a model of chromatin architecture in the cell. Chromosome Res., 19, 37–51.

37. Sazer, S. and Schiessel, H. (2018) The biology and polymer physics underlying large-scale chromosome organization. Traffic, 19, 87–104.

38. Rosa, A. (2022) The Physical Behavior of Interphase Chromosomes: Polymer Theory and Coarse-Grain Computer Simulations. Methods Mol. Biol., 2301, 235–258.

39. Kaplan, N. and Dekker, J. (2013) High-throughput genome scaffolding from in vivo DNA interaction frequency. Nat. Biotechnol., 31, 1143–1147.

40. Oddes, S., Zelig, A. and Kaplan, N. (2018) Three invariant Hi-C interaction patterns: Applications to genome assembly. Methods, 10.1016/j.ymeth.2018.04.013.

41. Burton, J.N., Adey, A., Patwardhan, R.P., Qiu, R., Kitzman, J.O. and Shendure, J. (2013) Chromosome-scale scaffolding of de novo genome assemblies based on chromatin interactions. Nat. Biotechnol., 31, 1119–25.

42. Marbouty, M., Cournac, A., Flot, J.-F., Marie-Nelly, H., Mozziconacci, J. and Koszul, R. (2014) Metagenomic chromosome conformation capture (meta3C) unveils the diversity of chromosome organization in microorganisms. Elife, 3.

43. Burton, J.N., Liachko, I., Dunham, M.J. and Shendure, J. (2014) Species-level deconvolution of metagenome assemblies with Hi-C-based contact probability maps. G3, 4, 1339–46.

44. Selvaraj, S., R Dixon, J., Bansal, V. and Ren, B. (2013) Whole-genome haplotype reconstruction using proximity-ligation and shotgun sequencing. Nat. Biotechnol., 31, 1111–8.

45. Rao, S.S.P., Huntley, M.H., Durand, N.C., Stamenova, E.K., Bochkov, I.D., Robinson, J.T., Sanborn, A.L., Machol, I., Omer, A.D., Lander, E.S., et al. (2014) A 3D map of the human genome at kilobase resolution reveals principles of chromatin looping. Cell, 159, 1665–80.

46. Sexton, T., Yaffe, E., Kenigsberg, E., Bantignies, F., Leblanc, B., Hoichman, M., Parrinello, H., Tanay, A. and Cavalli, G. (2012) Three-dimensional folding and functional organization principles of the Drosophila genome. Cell, 148, 458–72.

47. Dixon, J.R., Selvaraj, S., Yue, F., Kim, A., Li, Y., Shen, Y., Hu, M., Liu, J.S. and Ren, B. (2012) Topological domains in mammalian genomes identified by analysis of chromatin interactions. Nature, 485, 376–380.

48. Nora, E.P., Lajoie, B.R., Schulz, E.G., Giorgetti, L., Okamoto, I., Servant, N., Piolot, T., Van Berkum, N.L., Meisig, J., Sedat, J., et al. (2012) Spatial partitioning of the regulatory landscape of the X-inactivation centre. Nature, 485, 381–385.

49. Fudenberg, G., Imakaev, M., Lu, C., Goloborodko, A., Abdennur, N. and Mirny, L.A. (2016) Formation of Chromosomal Domains by Loop Extrusion. Cell Rep., 15, 2038–2049.

50. Sanborn, A.L., Rao, S.S.P., Huang, S.-C., Durand, N.C., Huntley, M.H., Jewett, A.I., Bochkov, I.D., Chinnappan, D., Cutkosky, A., Li, J., et al. (2015) Chromatin extrusion explains key features of loop and domain formation in wild-type and engineered genomes. Proc. Natl. Acad. Sci., 10.1073/pnas.1518552112.

51. Fudenberg, G., Abdennur, N., Imakaev, M., Goloborodko, A. and Mirny, L.A. (2017) Emerging Evidence of Chromosome Folding by Loop Extrusion. Cold Spring Harb. Symp. Quant. Biol., 82, 45–55.

52. Nora, E.P., Goloborodko, A., Valton, A.L., Gibcus, J.H., Uebersohn, A., Abdennur, N., Dekker, J., Mirny, L.A. and Bruneau, B.G. (2017) Targeted Degradation of CTCF Decouples Local Insulation of Chromosome Domains from Genomic Compartmentalization. Cell, 169, 930-944.e22.

53. Rao, S.S.P., Huang, S.C., Glenn St Hilaire, B., Engreitz, J.M., Perez, E.M., Kieffer-Kwon, K.R., Sanborn, A.L., Johnstone, S.E., Bascom, G.D., Bochkov, I.D., et al. (2017) Cohesin Loss Eliminates All Loop Domains. Cell, 171, 305-320.e24.

54. Imakaev, M., Fudenberg, G., McCord, R.P., Naumova, N., Goloborodko, A., Lajoie, B.R., Dekker, J. and Mirny, L.A. (2012) Iterative correction of Hi-C data reveals hallmarks of chromosome organization. Nat. Methods, 9, 999–1003.

55. Yaffe, E. and Tanay, A. (2011) Probabilistic modeling of Hi-C contact maps eliminates systematic biases to characterize global chromosomal architecture. Nat. Genet., 43, 1059–1065.

56. Nichols, M.H. and Corces, V.G. (2021) Principles of 3D compartmentalization of the human genome. Cell Rep., 35, 109330.

57. Zheng, X. and Zheng, Y. (2018) CscoreTool: Fast Hi-C compartment analysis at high resolution. Bioinformatics, 34, 1568–1570.

58. Di Pierro, M., Cheng, R.R., Aiden, E.L., Wolynes, P.G. and Onuchic, J.N. (2017) De novo prediction of human chromosome structures: Epigenetic marking patterns encode genome architecture. Proc. Natl. Acad. Sci. U. S. A., 114, 12126–12131.

59. Di Pierro, M., Zhang, B., Aiden, E.L., Wolynes, P.G. and Onuchic, J.N. (2016) Transferable model for chromosome architecture. Proc. Natl. Acad. Sci. U. S. A., 113, 12168–12173.

60. Fortin, J.P. and Hansen, K.D. (2015) Reconstructing A/B compartments as revealed by Hi-C using long-range correlations in epigenetic data. Genome Biol., 16, 1–23.

61. Xiong, K. and Ma, J. (2019) Revealing Hi-C subcompartments by imputing inter-chromosomal chromatin interactions. Nat. Commun., 10, 1–12.

62. Esposito, A., Bianco, S., Chiariello, A.M., Abraham, A., Fiorillo, L., Conte, M., Campanile, R. and Nicodemi, M. (2022) Polymer physics reveals a combinatorial code linking 3D chromatin architecture to 1D chromatin states. Cell Rep., 38.

63. Zhou, J. (2022) Sequence-based modeling of three-dimensional genome architecture from kilobase to chromosome scale. Nat. Genet., 54, 725–734.

64. Jost, D., Carrivain, P., Cavalli, G. and Vaillant, C. (2014) Modeling epigenome folding: formation and dynamics of topologically associated chromatin domains. Nucleic Acids Res., 42, 9553–61.

65. Jost, D. (2022) Polymer Modeling of 3D Epigenome Folding: Application to Drosophila. Methods Mol. Biol., 2301, 293–305.

66. Di Stefano, M., Nützmann, H.W., Marti-Renom, M.A. and Jost, D. (2021) Polymer modelling unveils the roles of heterochromatin and nucleolar organizing regions in shaping 3D genome organization in Arabidopsis thaliana. Nucleic Acids Res., 49, 1840–1858.

67. Olarte-Plata, J.D., Haddad, N., Vaillant, C. and Jost, D. (2016) The folding landscape of the epigenome. Phys. Biol., 13.

68. Nuebler, J., Fudenberg, G., Imakaev, M., Abdennur, N. and Mirny, L.A. (2018) Chromatin organization by an interplay of loop extrusion and compartmental segregation. Proc. Natl. Acad. Sci., 115, E6697–E6706.

69. Falk, M., Feodorova, Y., Naumova, N., Imakaev, M., Lajoie, B.R., Leonhardt, H., Joffe, B., Dekker, J., Fudenberg, G., Solovei, I., et al. (2019) Heterochromatin drives compartmentalization of inverted and conventional nuclei. Nature, 570, 395–399.

70. Bianco, S., Lupiáñez, D.G., Chiariello, A.M., Annunziatella, C., Kraft, K., Schöpflin, R., Wittler, L., Andrey, G., Vingron, M., Pombo, A., et al. (2018) Polymer physics predicts the effects of structural variants on chromatin architecture. Nat. Genet., 50, 662–667.

71. Barbieri, M., Chotalia, M., Fraser, J., Lavitas, L.M., Dostie, J., Pombo, A. and Nicodemi, M. (2012) Complexity of chromatin folding is captured by the strings and binders switch model. Proc Natl Acad Sci U S A, 109, 16173–16178.

72. Chiariello, A.M., Annunziatella, C., Bianco, S., Esposito, A. and Nicodemi, M. (2016) Polymer physics of chromosome large-scale 3D organisation. Sci. Rep., 6, 1–8.

73. Schwarzer, W., Abdennur, N., Goloborodko, A., Pekowska, A., Fudenberg, G., Loe-Mie, Y., Fonseca, N.A., Huber, W., H Haering, C., Mirny, L., et al. (2017) Two independent modes of chromatin organization revealed by cohesin removal. Nature, 551, 51–56.

74. Zhang, Y., An, L., Xu, J., Zhang, B., Zheng, W.J., Hu, M., Tang, J. and Yue, F. (2018) Enhancing Hi-C data resolution with deep convolutional neural network HiCPlus. Nat. Commun., 9, 1–9.

75. Li, Z. and Dai, Z. (2020) SRHiC: A Deep Learning Model to Enhance the Resolution of Hi-C Data. Front. Genet., 11, 1–9.

76. Hong, H., Jiang, S., Li, H., Du, G., Sun, Y., Tao, H., Quan, C., Zhao, C., Li, R., Li, W., et al. (2020) DeepHic: A generative adversarial network for enhancing Hi-C data resolution. PLoS Comput. Biol., 16, 1–25.

77. Liu, T. and Wang, Z. (2019) HiCNN: A very deep convolutional neural network to better enhance the resolution of Hi-C data. Bioinformatics, 35, 4222–4228.

78. Parada, L.A., McQueen, P.G. and Misteli, T. (2004) Tissue-specific spatial organization of genomes. Genome Biol., 5, R44.

79. Ganai, N., Sengupta, S. and Menon, G.I. (2014) Chromosome positioning from activity-based segregation. Nucleic Acids Res., 42, 4145–4159.

80. Bernstein, B.E., Birney, E., Dunham, I., Green, E.D., Gunter, C. and Snyder, M. (2012) An integrated encyclopedia of DNA elements in the human genome. Nature, 489, 57–74.

81. Kim, H.J., Yardımcı, G.G., Bonora, G., Ramani, V., Liu, J., Qiu, R., Lee, C., Hesson, J., Ware, C.B., Shendure, J., et al. (2020) Capturing cell type-specific chromatin compartment patterns by applying topic modeling to single-cell Hi-C data. PLoS Comput. Biol., 16, 1–19.

82. Ernst, J. and Kellis, M. (2012) ChromHMM: automating chromatin-state discovery and characterization. Nat. Methods, 9, 215–6.

83. Filion, G.J., van Bemmel, J.G., Braunschweig, U., Talhout, W., Kind, J., Ward, L.D., Brugman, W., de Castro, I.J., Kerkhoven, R.M., Bussemaker, H.J., et al. (2010) Systematic Protein Location Mapping Reveals Five Principal Chromatin Types in Drosophila Cells. Cell, 143, 212–224.

84. Reiff, S.B., Schroeder, A.J., Kırlı, K., Cosolo, A., Bakker, C., Lee, S., Veit, A.D., Balashov, A.K., Vitzthum, C., Ronchetti, W., et al. (2022) The 4D Nucleome Data Portal as a resource for searching and visualizing curated nucleomics data. Nat. Commun., 13, 1–11.

85. Abdennur, N. and Mirny, L.A. (2020) Cooler: Scalable storage for Hi-C data and other genomically labeled arrays. Bioinformatics, 36, 311–316.

86. Pedregosa, F., Weiss, R. and Brucher, M. (2011) Scikit-learn□: Machine Learning in Python. J. Mach. Learn. Res., 12, 2825–2830.

87. Behnel, S., Bradshaw, R., Citro, C., Dalcin, L., Seljebotn, D.S. and Smith, K. (2011) Cython: The Best of Both Worlds. Comput. Sci. Eng., 13, 31–39.

88. Harris, C.R., Millman, K.J., van der Walt, S.J., Gommers, R., Virtanen, P., Cournapeau, D., Wieser, E., Taylor, J., Berg, S., Smith, N.J., et al. (2020) Array programming with NumPy. Nature, 585, 357–362.

